## Supplementary Information for "Synchrony, oscillations, and phase relationships in collective neuronal activity: a highly comparative overview of methods"

### Supporting information

**Table S1. Number of spike trains for each synthetic spike train family and biological dataset.**

| name | $N_{\mathrm{win}}^{\mathrm{tot}}$ | $N_{\mathrm{win}}^{\mathrm{rej}}$, core set | $N_{\mathrm{win}}^{\mathrm{rej}}$, extended set | $N_{\mathrm{win}}^{\mathrm{rej}}/N_{\mathrm{win}}^{\mathrm{tot}}$, core set | $N_{\mathrm{win}}^{\mathrm{rej}}/N_{\mathrm{win}}^{\mathrm{tot}}$, extended set |
|---|---|---|---|---|---|
| **Synthetic spike train families** | | | | | |
| single-scale | 450 | 21 | 52 | 4.67% | 11.56% |
| dual-scale | 450 | 124 | 190 | 27.56% | 42.22% |
| **Biological datasets** | | | | | |
| rat A1 | 280 | 6 | 8 | 2.14% | 2.86% |
| mouse CA1 | 800 | 219 | 296 | 27.37% | 37% |
| monkey V1 | 294 | 255 | 257 | 86.73% | 87.41% |

$N_{\mathrm{win}}^{\mathrm{tot}}$, total number of time windows; $N_{\mathrm{win}}^{\mathrm{rej}}$, number of time windows with at least one excluded MSTM.

**Table S2. Number of time windows in each class for each recording.**

| recording | $N_{\mathrm{win}}^{\mathrm{wake}}$ | $N_{\mathrm{win}}^{\mathrm{NREM}}$ | $N_{\mathrm{win}}^{\mathrm{wake,non-rej}}$ | $N_{\mathrm{win}}^{\mathrm{NREM,non-rej}}$ |
|---|---|---|---|---|
| M1R1 | 58 | 55 | 19 | 29 |
| M1R2 | 61 | 28 | 33 | 23 |
| M2R2 | 22 | 13 | 12 | 11 |
| M4R1 | 33 | 71 | 15 | 33 |

Number of time windows in each class for each recording of the mouse dataset (Formozov et al., 2022b) used for the brain state decoding analysis in section "Brain state classification from spike train measures". $N_{\mathrm{win}}^{\mathrm{wake,non-rej}}$ and $N_{\mathrm{win}}^{\mathrm{NREM,non-rej}}$ indicate the number of time windows classified as wake and NREM, respectively, where no MSTM was rejected.

**Figure S1. Assessment of synchrony on synthetic spike trains.** Assessment of the level of synchrony on single-scale (left) and dual-scale (right) synthetic spike trains, considering multiple MSTMs. Left: Synchrony is plotted as a function of the modulation amplitude $m$ for average firing rates $r_0$ and population rates $f_0$ varying in the grid [4,12,36] Hz × [4,12,36] Hz. Results for different average firing rates $r_0$ are plotted in separate columns. Right: Synchrony is plotted as a function of the population width $\Sigma$ for population rates $f_0$ and spike deletion probability $p_{\mathrm{fail}}$ varying in the grid [4,12,36] Hz × [0.8,0.4,0]. Results for different spike deletion probabilities $p_{\mathrm{fail}}$ are plotted in separate columns. In all panels, the population rate $f_0$ varies in the grid [4,12,36] Hz and is color-coded, with warmer colors indicating higher $f_0$. Solid lines: pseudo-rhythmic spike trains; dash lines: non-rhythmic spike trains.

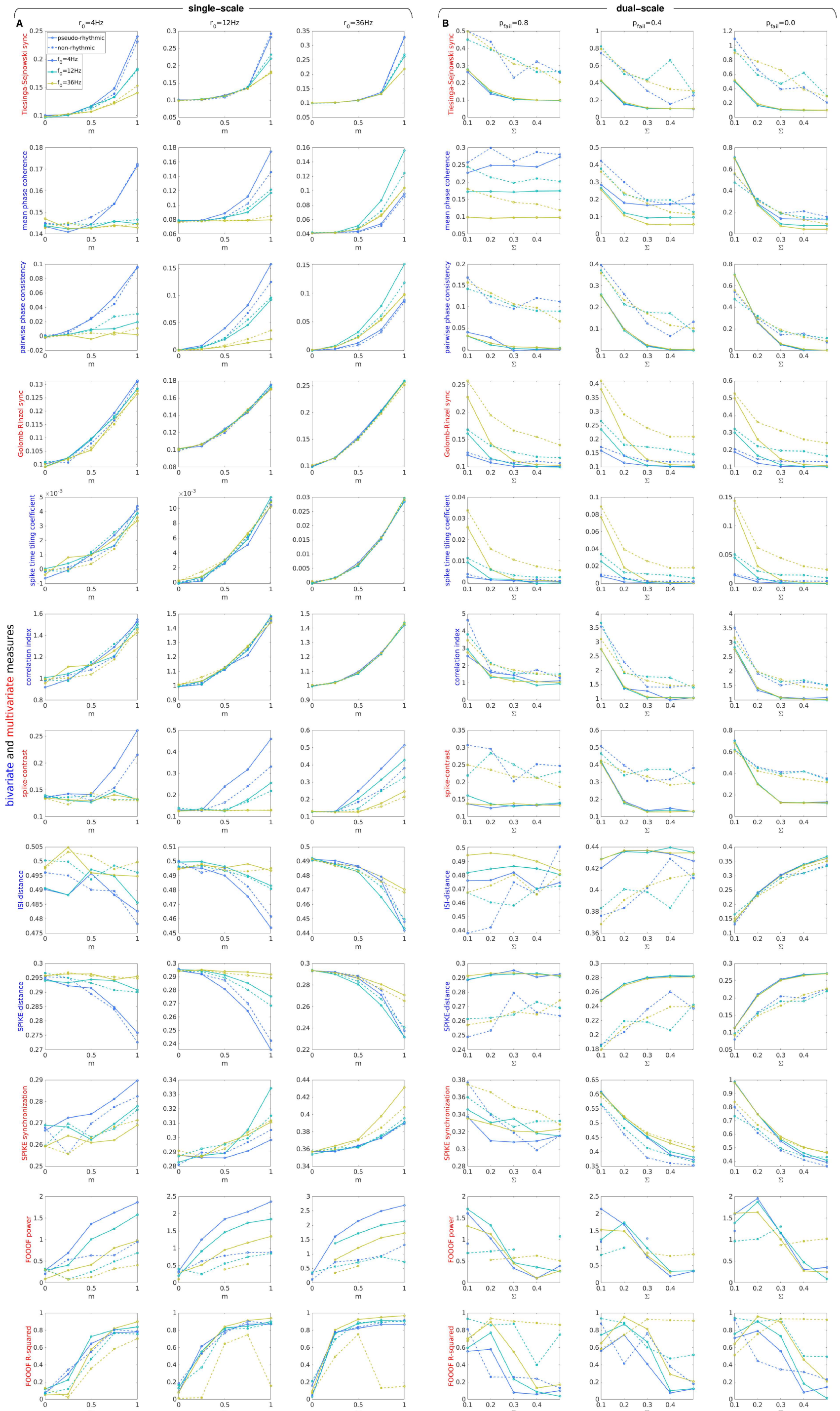
single-scale
dual-scale
A
B
$r_0$=4Hz
$r_0$=12Hz
$r_0$=36Hz
$p_{fail}$=0.8
$p_{fail}$=0.4
$p_{fail}$=0.0
pseudo-rhythmic
non-rhythmic
$f_0$=4Hz
$f_0$=12Hz
$f_0$=36Hz
bivariate and multivariate measures
Tiesinga-Sejnowski sync
mean phase coherence
pairwise phase consistency
Golomb-Rinzel sync
spike time tiling coefficient
correlation index
spike-contrast
ISI-distance
SPIKE-distance
SPIKE synchronization
FOOOF power
FOOOF R-squared
m
Σ

**Figure S2. Hierarchical clustering of measures, considering multiple timescales for the timescale-specific measures.** A: Dendrogram showing the distances between MSTMs as a hierarchical cluster tree. Measures are sorted along the $x$ axis in increasing order of the dendrogrammatic distance of the first nonsingleton cluster they are grouped in. B: Similarity matrix showing the absolute value of the Spearman correlation coefficient between each pair of MSTMs. Measure labels are color-coded to indicate measure type (green: univariate; blue: bivariate; red: multivariate). Numbers displayed at the end of timescale-dependent measures indicate the timescale in milliseconds. For FOOOF measures, `1p` (`2p`) refers to the single-peak (dual-peak) model; in the dual-peak model, the number at the end of the parameter name indicates the corresponding peak.

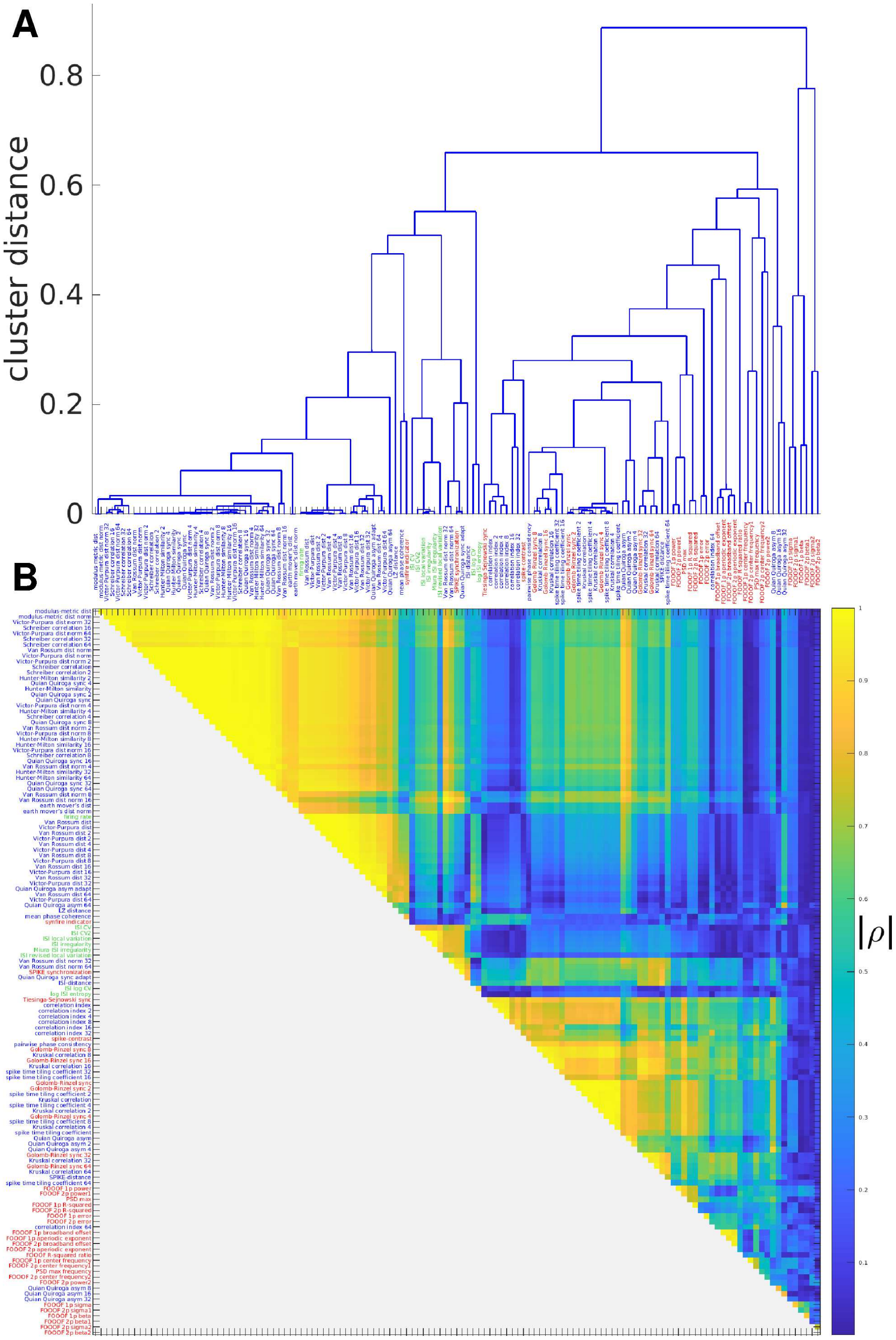
A
cluster distance
0.8
0.6
0.4
0.2
0
B
|ρ|

**Figure S3. Effects of time window length and number of neurons for a subset of selected measures.** Effects of sample size variations in time (A) or space (B) for a subset of selected MSTMs. A: For each MSTM, the mean value across windows is plotted as a function of window length. Shaded areas indicate the SD across windows. Different columns correspond to different spike train families and generative parameter values: single-scale pseudo-rhythmic spike train with $r_0 = f_0 = 12\,\mathrm{Hz}$, $m = 0.5$ (non-sequential, left; sequential with $D_c = 0.2$, center-left); dual-scale pseudo-rhythmic spike train with $f_0 = 12\,\mathrm{Hz}$, $\Sigma = 0.2$, $p_{\mathrm{fail}} = 0.4$, (non-sequential, center-right; sequential with $D_c = 0.2$, right). MPC and PPC could not be estimated at the shortest time window length considered due to insufficient number of spikes. B: As in (A), but sample size varies in space instead of time.

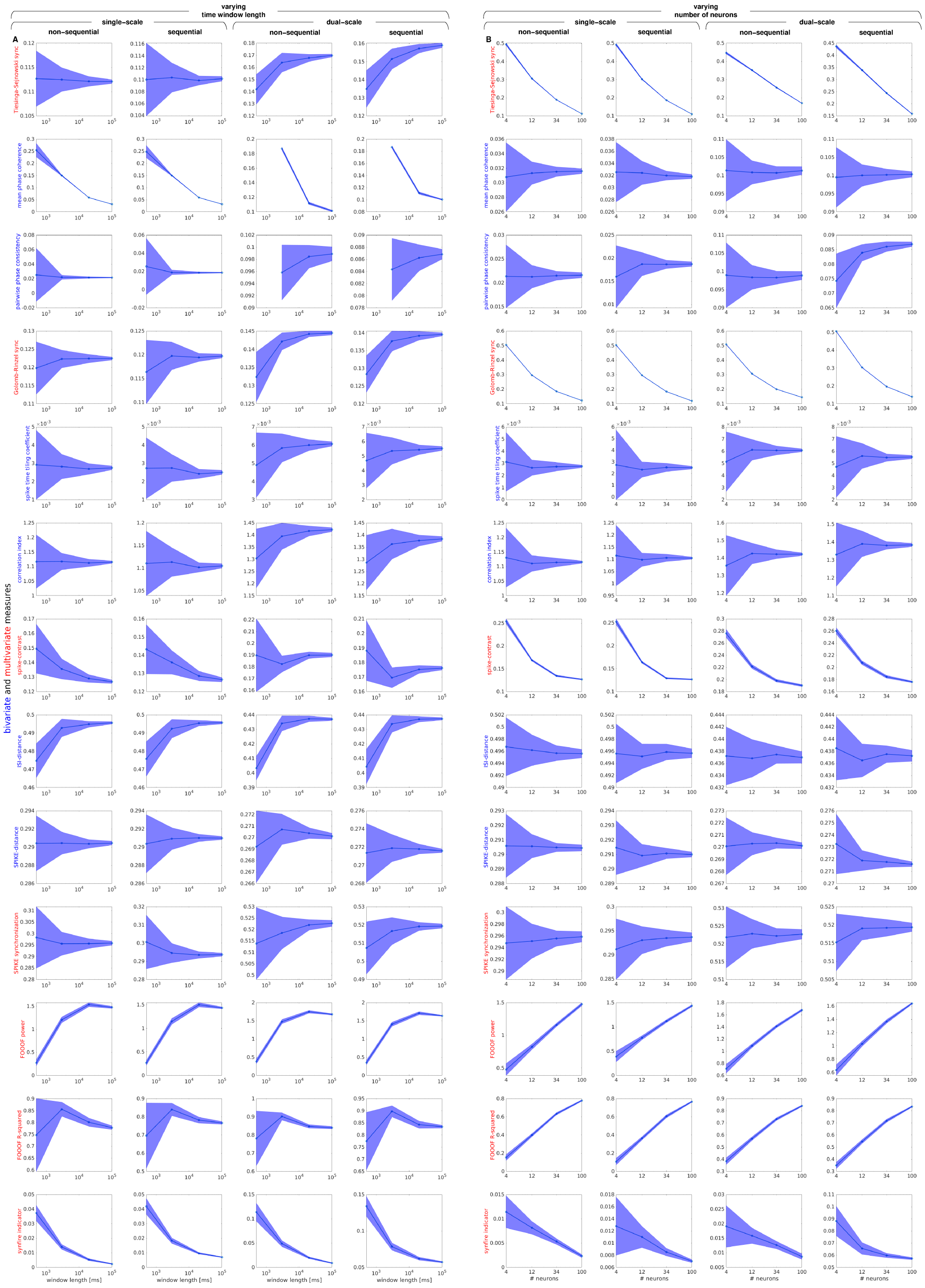

varying
time window length
single-scale
dual-scale
non-sequential
sequential
varying
number of neurons
A
B
bivariate and multivariate measures
Tiesinga-Sejnowski sync
mean phase coherence
pairwise phase consistency
Golomb-Rinzel sync
spike time tiling coefficient
correlation index
spike-contrast
ISI-distance
SPIKE-distance
SPIKE synchronization
FOOOF power
FOOOF R-squared
synfire indicator
window length [ms]
### neurons

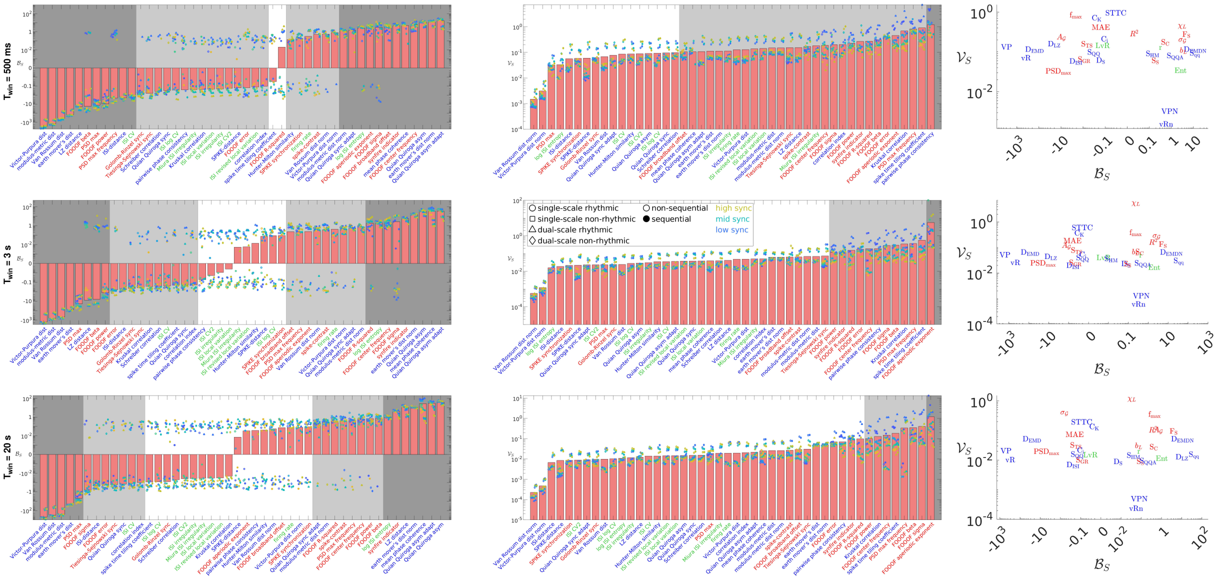

**Figure S4. Bias and variability resulting from a finite time window length at different timescales.** The same information plotted in Fig. 6A is replotted separately for each value of the time window length $T_{\mathrm{win}}$: results corresponding to short ($T_{\mathrm{win}} = 500\,\mathrm{ms}$), intermediate ($T_{\mathrm{win}} = 3\,\mathrm{s}$) and long ($T_{\mathrm{win}} = 20\,\mathrm{s}$) time windows are plotted in the top, middle, and bottom row, respectively.

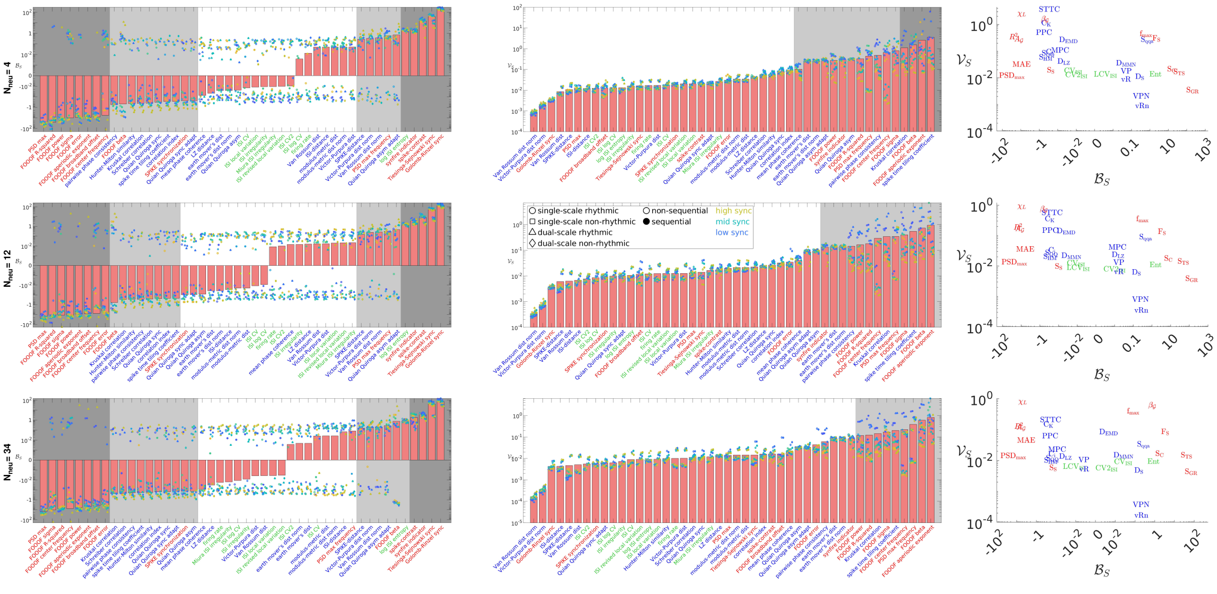

**Figure S5. Bias and variability resulting from a finite number of neurons at different spatial scales.** The same information plotted in Fig. 6B is replotted separately for each value of the number of neurons $N_{\mathrm{neu}}$: results corresponding to a low ($N_{\mathrm{neu}} = 4$), intermediate ($N_{\mathrm{neu}} = 12$) and high ($N_{\mathrm{neu}} = 34$) number of neurons are plotted in the top, middle, and bottom row, respectively.

**Figure S6. Visualizing collective dynamic coordination in spiking activity through the lens of the highly comparative approach.** As in Fig. 7, for the mouse (A) and monkey (B) datasets.

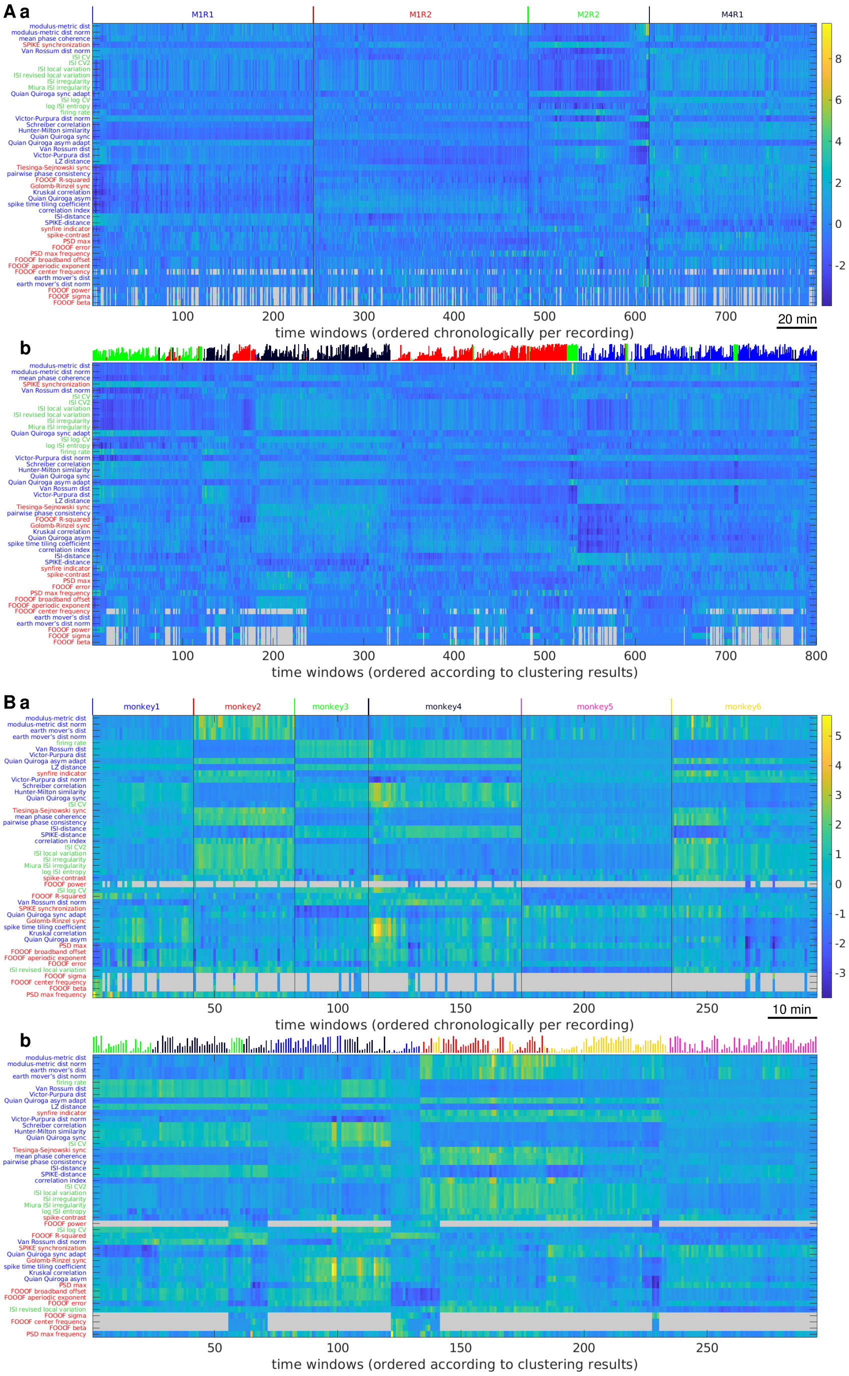


**Figure S7. Spiking activity exhibits structured variability that is distinctive of each recording.** As in Fig. 8, for the mouse (A) and monkey (B) datasets.

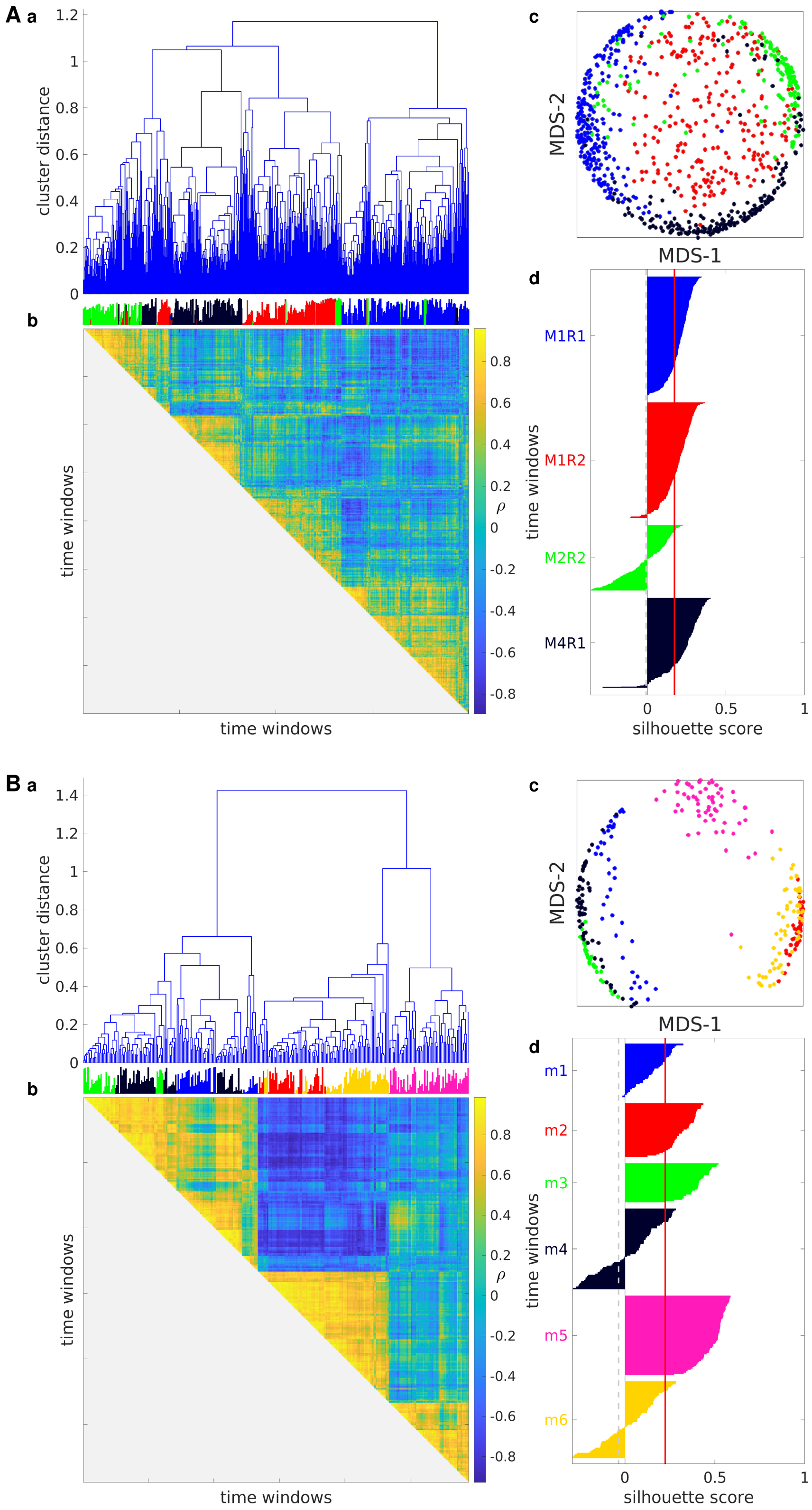
A a
cluster distance
b
time windows
time windows
ρ
c
MDS-2
MDS-1
d
M1R1
M1R2
M2R2
M4R1
time windows
silhouette score
B a
cluster distance
b
time windows
time windows
ρ
c
MDS-2
MDS-1
d
m1
m2
m3
m4
m5
m6
time windows
silhouette score

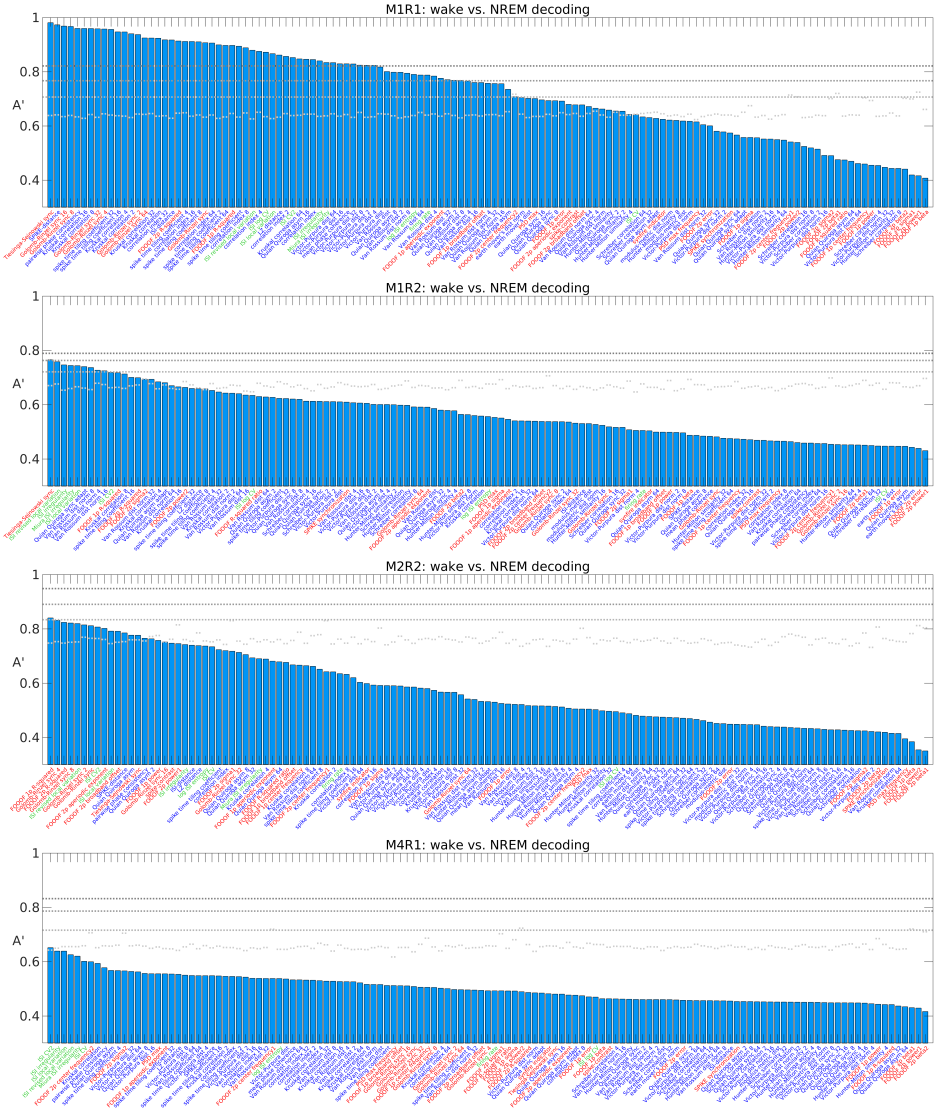


**Figure S8. Univariate decoding results for each individual recording and for each MSTM of the extended set of 131 MSTMs.** As in Fig. 9A, for each individual recording and for each MSTM of the extended set of 131 MSTMs.

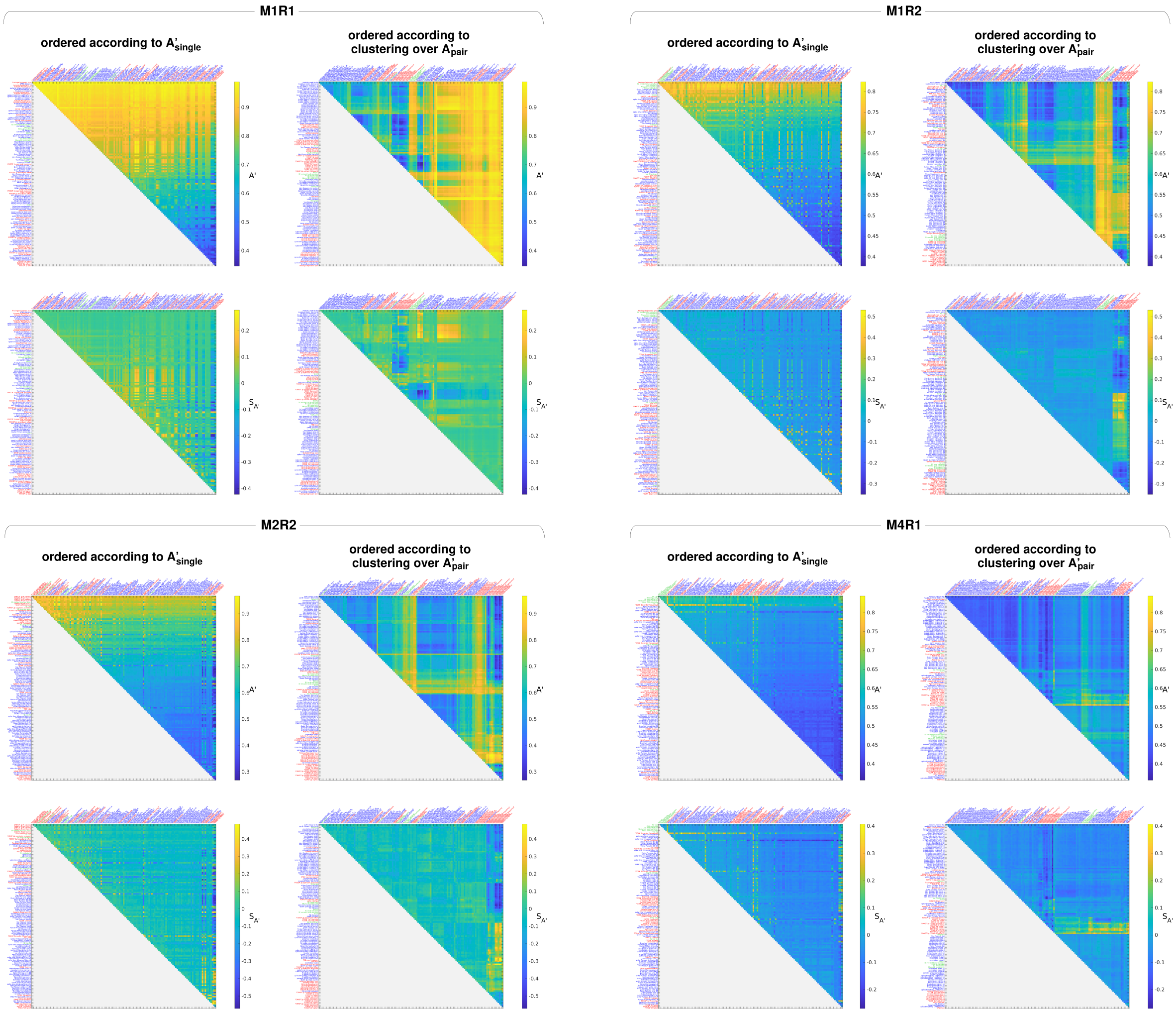


**Figure S9. Bivariate decoding results for each individual recording and for each MSTM of the extended set of 131 MSTMs.** As in Fig. 10, for each individual recording and for each MSTM of the extended set of 131 MSTMs. The same results are shown with the MSTMs ordered according to decreasing univariate decoding accuracy (left column), and with the MSTMs ordered according to the results of a hierarchical agglomerative clustering analysis over bivariate decoding accuracies (right column), hence displaying nearby MSTMs that result in similar patterns of decoding accuracies when combined with every other MSTMs.

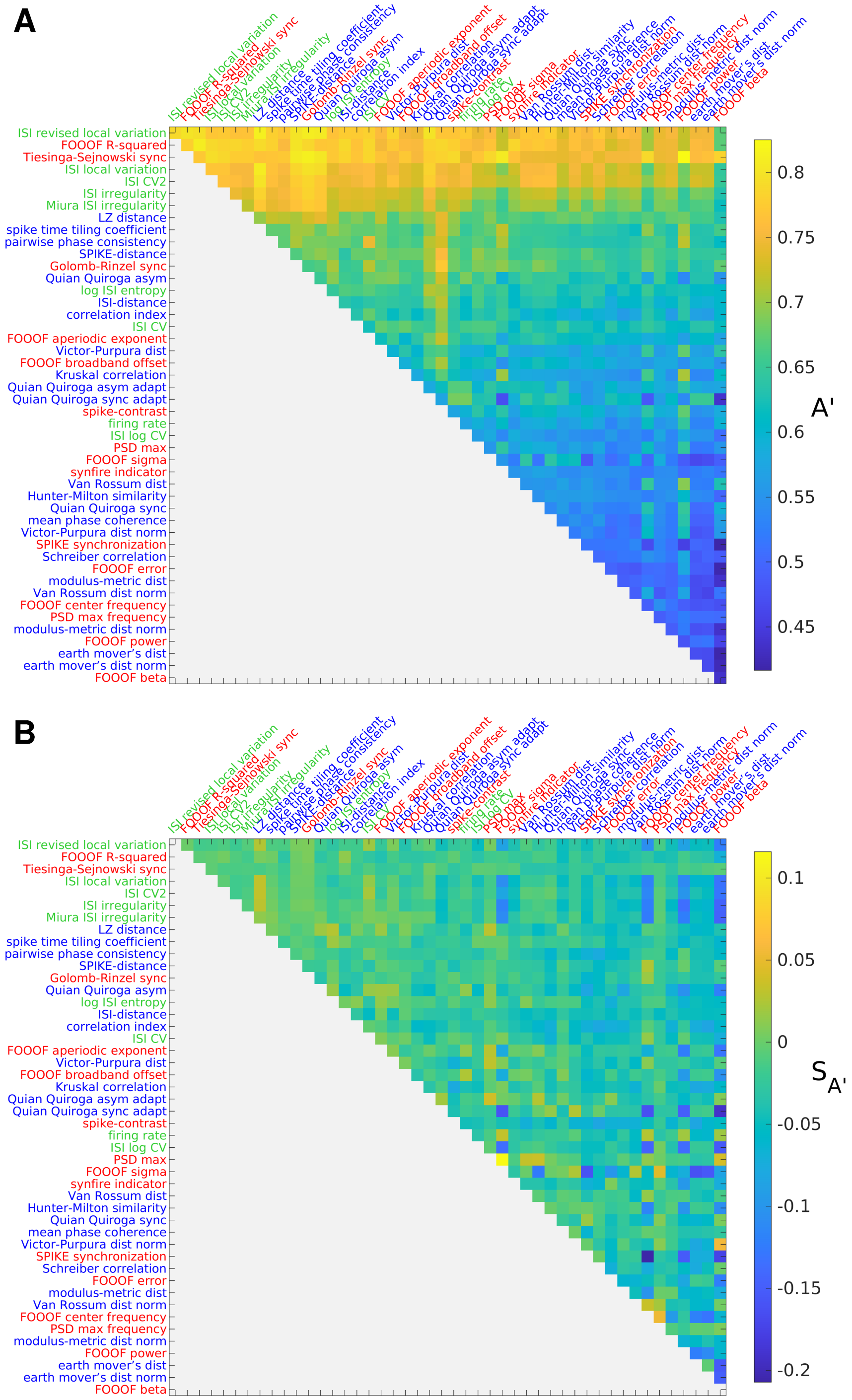


**Figure S10. Discriminating wake vs. NREM sleep from spike train measure pairs: median results across recordings.** As in Fig. 10, but median results across recordings are shown.
